# Transcription factor LSF facilitiates lysine methylation of α-tubulin by microtubule-associated SET8

**DOI:** 10.1101/665984

**Authors:** Hang Gyeong Chin, Pierre Olivier Estève, Cristian Ruse, Jiyoung Lee, Scott E. Schaus, Sriharsa Pradhan, Ulla Hansen

## Abstract

Microtubules are critical for mitosis, cell motility, and protein and organelle transport, and are a validated target for anticancer drugs. However, tubulin regulation and recruitment in these cellular processes is less understood. Post-translational modifications of tubulin are proposed to regulate microtubule functions and dynamics. Although many such modifications have been investigated, tubulin methylations and enzymes responsible for methylation have only recently begun to be described. Here we report that N-lysine methyl transferase KMT5A (SET8/PR-Set7), which methylates histone H4K20, also methylates α-tubulin. Furthermore, the transcription factor LSF binds both tubulin and SET8, and enhances α-tubulin methylation *in vitro*, countered by FQI1, a specific small molecule inhibitor of LSF. Thus, the three proteins SET8, LSF, and tubulin, all essential for mitotic progression, interact with each other. Overall, these results point to dual functions for both SET8 and LSF not only in chromatin regulation, but also for cytoskeletal modification.

## Introduction

Microtubules (MTs), the polymerized heterodimers of α-tubulin and β-tubulin, are major cytoskeletal components that play important roles in key cellular processes such as structural support, localization of organelles, and chromosome segregation (Janke, 2014;Verhey and Gaertig, 2007). A number of post-translational modifications (PTMs) of tubulins have been reported, which contribute to the functional diversity of MTs and affect MT dynamics and organization (Song and Brady, 2015). This led to the hypothesis of a tubulin code (Verhey *et al.*, 2007), where tubulin modifications specify biological outcomes through changes in higher-order microtubule structure by recruiting and interacting with effector proteins. Notably, tubulin methylation has been less studied than other types of tubulin modification, such as tyrosination, glutamylation, glycylation, acetylation, and phosphorylation, although in the parallel histone code hypothesis, methylation is the most common and well-understood modification.

SET8/PR-Set7 is a N-lysine methyltransferase responsible for the monomethylation of both histone and non-histone proteins in higher eukaryotes (Dillon *et al.*, 2005). It is functionally characterized as a histone H4 lysine 20-specific monomethyltransferase (Fang *et al.*, 2002); this modification is a specific mark for transcriptional repression and is also enriched during mitosis (Nishioka *et al.*, 2002;Rice *et al.*, 2002). SET8 is required for cell proliferation, chromosome condensation, and cytokinesis, since deletion or RNAi mediated depletion of the enzyme impairs all these functions. Previous findings, in particular, suggested that SET8 and H4K20me1 are required for mitotic entry (Wu and Rice, 2011). SET8 also mediates monomethylation of other substrates, including p53, which results in repression of p53 target genes (Shi *et al.*, 2007). However, how H4K20me1 is regulated and how it functions to promote cell cycle progression remains an open question, including the possibility that other non-histone substrates may be involved.

LSF, previously characterized widely as a transcription factor, is an oncogene in hepatocellular carcinoma, being signficantly overexpressed in hepatocellular carcinoma cell lines and patient samples (Fan *et al.*, 2011;Gu *et al.*, 2015;Kim *et al.*, 2017;Seol *et al.*, 2016;Yoo *et al.*, 2010;Zhang *et al.*, 2017), as well as in other cancer types (Kotarba *et al.*, 2018). LSF is also generally required for cell cycle progression and cell survival (Hansen *et al.*, 2009;Powell *et al.*, 2000;Rajasekaran *et al.*, 2015). Initially, LSF was described as a regulator of G1/S progression (Powell *et al.*, 2000) and essential for inducing expression of the gene encoding thymidylate synthase (TYMS) in late G1. The additional involvement of LSF in mitosis resulted from characterization of the effects of Factor Quinolinone Inhibitor 1 (FQI1), a specific small molecule inhibitor of LSF (Rajasekaran *et al.*, 2015). FQI1 not only abrogates the DNA-binding and corresponding transcriptional activities of LSF (Rajasekaran *et al.*, 2015), but also specific LSF-protein interactions (Chin *et al.*, 2016). Finally, FQI1 inhibits growth of hepatocellular carcinoma tumors in multiple mouse models, and causes cell death via mitotic defects in hepatocellular carcinoma cell lines (Grant *et al.*, 2012;Rajasekaran *et al.*, 2015).

In this study, we demonstrate that these three regulators of mitosis, SET8, LSF, and tubulin, all interact with each other both *in vitro* and within cells. Furthermore, we demonstrate that SET8 is a microtubule-associated methyltransferase that methylates lysines on α-tubulin. In parallel to how transcription factors stimulate histone modification by interacting both with the chromatin writers and the DNA, LSF stimulates methylation of tubulin by SET8. These results suggest that LSF and SET8 have biological implications beyond gene transcription and histone methylation, respectively.

## Results

### SET8 directly interacts with tubulin

Analysis of two commercial tubulin preparations (>97% and >99 % pure, respectively) by mass spectroscopy identified anticipated associated proteins (e.g. MAP1, MAP2), but surprisingly also peptides covering SET8 (Appendix Fig S1A; Appendix Table S1). The presence of SET8 in these preparations was indicated by immunoblotting, although it is a minor component (Appendix Fig S1B). This raised the question of whether SET8 might target MT-related substrates. Indeed, initial overexpression of GFP-SET8 in a COS7 cell line indicated that the majority of GFP-SET8 was localized in the cytoplasm (Fig 1A). Upon screening for association with various cytoplasmic structural features by staining with relevant fluorescence dyes or antibodies along with GFP-SET8 expression, GST-SET8 significantly co-localized only with α-tubulin, indicating MTs. MT co-localization was observed at stages through the cell cycle (Fig 1A, Appendix Fig S1C). Most obvious was in G1 phase, when SET8 exhibited the same pattern as the filamentous tubulin distributed throughout the cytoplasm, emphasized by the yellow in the merged image. In S phase, as expected from previous studies, some SET8 was also nuclear (Fig 1A). Although SET8 is often considered to be a nuclear protein, localization of SET8 also in the cytoplasm of human cells has been documented by others, in a cell type-specific manner (Thul *et al.*, 2017). Furthermore, biochemical fractionation confirmed localization of endogenous SET8 in both the nucleus and the cytoplasm of human HEK293T cells (Fig 1B), which were used in subsequent experiments.

**Figure 1.**
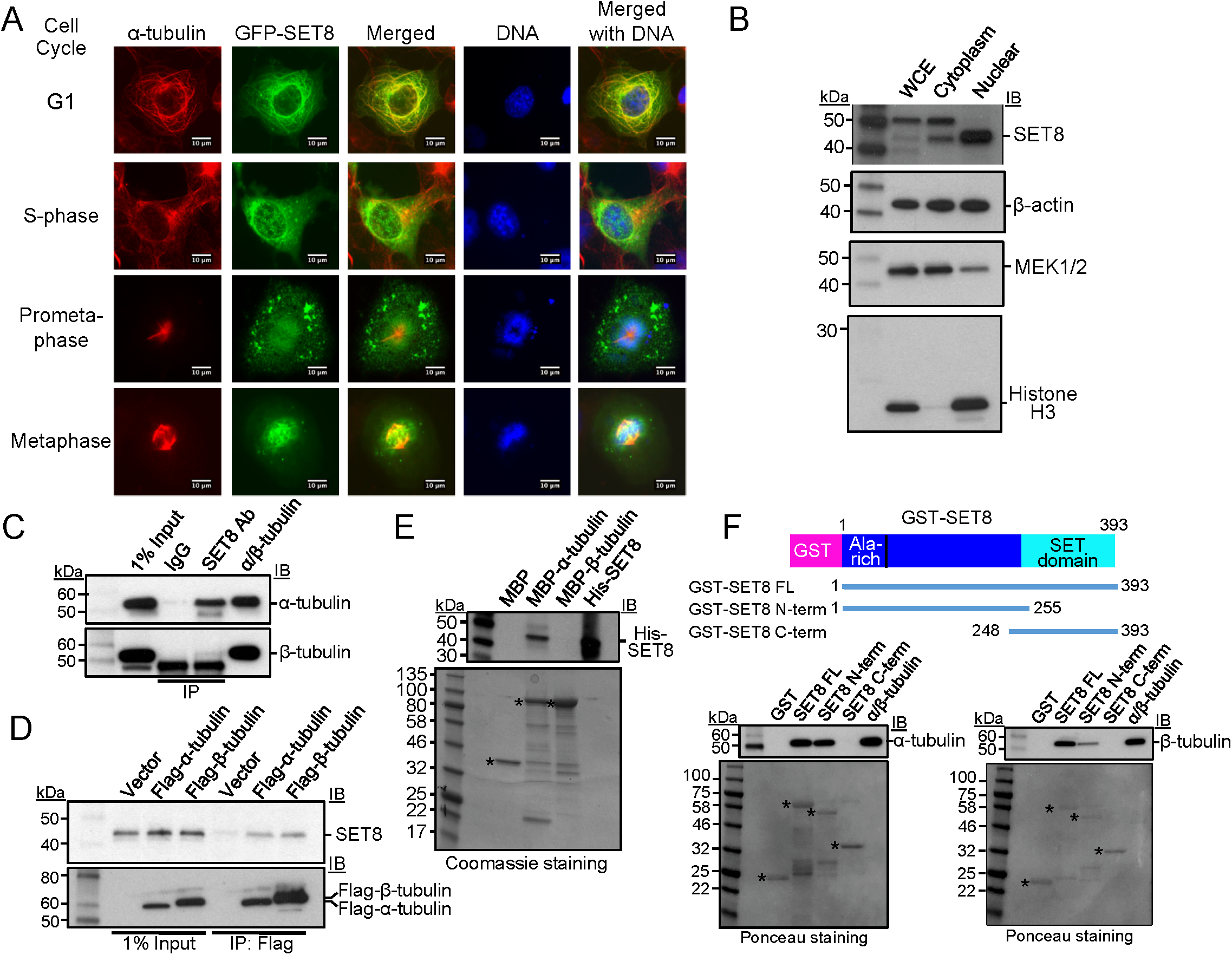
SET8 associates with tubulin in cells and directly interacts with α-tubulin *in vitro*. **A** Colocalization of SET8 and α-tubulin in COS7 cells. GFP-SET8 (green) was expressed in asynchronous cells, tubulin was detected with anti-α-tubulin antibody (red) and DNA with DAPI (blue). Yellow in the merged image indicates colocalization of SET8 and α-tubulin. Images are from cells identified as being in the indicated stages of cell cycle progression. Scale bars: 10 µm. **B** Endogenous SET8 is localized in both nucleus and cytoplasm in HEK293T cells. Whole cell extract (WCE), cytoplasmic, and nuclear fractions were separated and analyzed for the presence of SET8, β-actin (present in both), MEK1/2 (predominantly cytoplasmic marker), and histone H3 (nuclear marker). **C** Co-immunoprecipitation from HEK293T cells of endogenous tubulins with endogenous SET8, using SET8 antibody (Ab). Right lane: >99% pure tubulin (MP Biomedical, #0877112) as a positive control. Immunoprecipitates were analyzed using antibodies against the indicated proteins by immunoblotting (IB). **D** Co-immunoprecipitation from HEK293T cells of endogenous SET8 with transiently expressed Flag-tagged tubulins, as detected by immunoprecipitation with antibody against Flag. **E** MBP-pull down analysis of purified His-SET8 with MBP-α-tubulin, but not MBP-β-tubulin. Top: Immunoblot (IB) in which biotinylated molecular weight markers are visualized; bottom: Coomassie staining of the same gel (shown in grayscale) in which standard molecular weight markers are visualized. *Expected positions of migration of the MBP-proteins. **F** GST-pull down analysis of purified porcine brain tubulin to full-length or the indicated overlapping segments of SET8 fused to GST. Top: immunoblot (IB) in which biotinylated molecular weight markers are visualized; bottom: Ponceau staining of the same gels (shown in grayscale) in which standard molecular weight markers are visualized. *Expected positions of migration of the GST-proteins.

To determine whether endogenous cellular SET8 associates with tubulins, we immunoprecipitated protein complexes from HEK293T cell extracts. Using antibody against SET8, α-tubulin was also precipitated, as well as β-tubulin, although to a considerably lesser extent (Fig 1C). Conversely, upon expression of Flag-tagged α-tubulin or β-tubulin in the cells, endogenous SET8 co-immunoprecipitated with both, to roughly similar extents compared to the level of expression of the tagged tubulin (Fig 1D). As α- and β-tubulins stably heterodimerize in cells, *in vitro* experiments were required in order to determine whether either of these interactions was direct. To this end, purified recombinant proteins fusing maltose binding protein (MBP) to either α-tubulin (TUBA1A) or β-tubulin (TUBB) were individually tested for interactions with His-tagged SET8 purified from *E. coli*. SET8 directly interacted only with α-tubulin, but not with β-tubulin (Fig 1E). Conversely, recombinant proteins fusing glutathione S-transferase (GST) to either full-length, or the N- or C-terminal overlapping portions of human SET8 were tested for interactions *in vitro* with a purified tubulin preparation. The purified heterodimeric tubulin interacted only with the full-length and N-terminal portion of SET8, even though th C-terminal SET8 fusion protein was present at a higher level than the others (Fig 1F), indicating specificity of this interaction. Taken together, these data demonstrate that α-tubulin and SET8 directly interact with each other, whereas β-tubulin is only in a complex with SET8 in the presence of α-tubulin.

### SET8 methylates α-tubulin

SET8 was characterized historically as a nucleosomal H4K20 specific methyltransferase, and subsequently as a regulator of the non-histone protein p53. However, since SET8 bound strongly to α-tubulin, we tested whether tubulins could be a novel substrate of the enzyme. Purified mammalian α/β-tubulin was incubated with the cofactor S-adenosyl-L-[methyl-^3^H] methionine (AdoMet) and purified, recombinant GST-SET8. In the presence of both SET8 and AdoMet, radioactivity was incorporated into a protein band migrating at the position of α- and β-tubulins, in addition to a less pronounced automethylation of GST-SET8 (Fig 2A, lane 3), but no radioactive product was present at the position of α- and β-tubulins when either tubulin or SET8 was omitted from the reaction (Fig 2A, lanes 2 and 4). Interestingly, when histone H4 was also included in the reaction, the amount of tubulin modification was reduced (Fig 2A, lane 1), indicating that histone H4 strongly competed with tubulins for the methylation activity of SET8. Furthermore, histone H4 also competed with SET8 itself as a substrate, as shown by the significant reduction in SET8 automethylation in the presence of histone H4.

**Figure 2.**
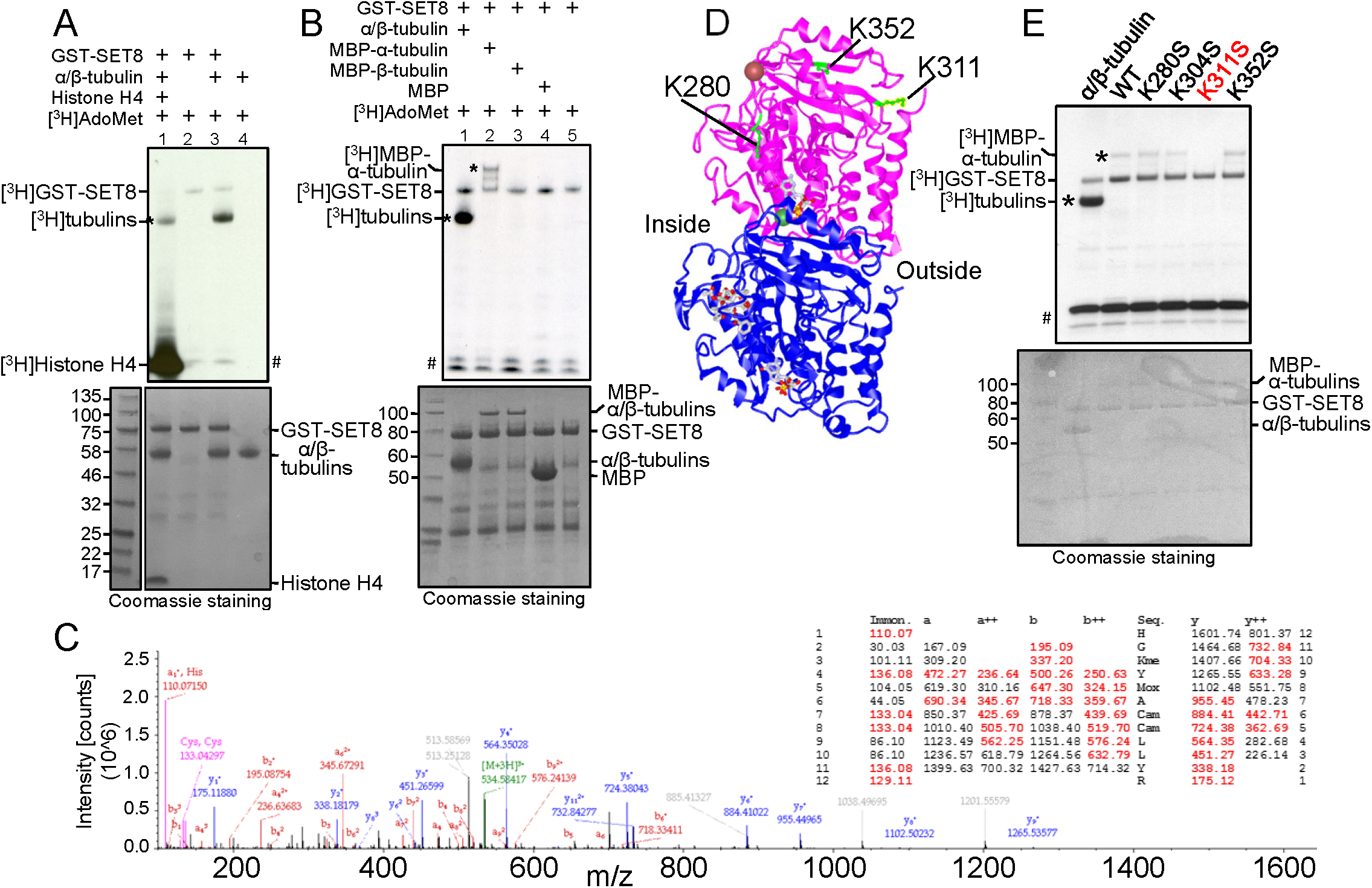
Histone methytransferase SET8 methylates α-tubulin at K311. **A** Purified porcine tubulin (rPeptide, # T-1201-1) is methylated by SET8. Lane 1: histone H4 (1 µg) was added in addition to tubulin as substrates. Top: autoradiogram of methyltransferase assays, showing methylation of tubulin (*), histone H4, and automethylation of GST-SET8. # indicates the migration of [^3^H]-labeled impurities, which migrated at a similar position to that of Histone H4. Bottom: Coomassie-staining of the same gel (shown in grayscale) indicating relative levels of the components in the reaction. **B** Recombinant human MBP-α-tubulin (*), but not MBP-β-tubulin, is methylated by SET8. Autoradiogram (top) and Coomassie-staining (bottom) are as described in **A**. Protein bands <50 kDa are from the purified GST-SET8 preparation, and are more evident in this experiment than in other reactions. **C** Mass spectrum and table of expected m/z of the peptide fragments (with observed fragments in red) confirming methylation on K311 of α-tubulin after incubation of purified tubulin with SET8. **D** The 3-dimensional structure of the α/β-tubulin heterodimer (PDB ID 1JFF; purple: α-tubulin; blue: β-tubulin), indicating positions of lysines in α-tubulin targeted by SET8 *in vitro* (green). Inside and outside surfaces of the MT structure are indicated. **E** Mutation solely of K311 in recombinant MBP-α-tubulin (K311S) substantially reduced methylation by GST-SET8 *in vitro*. Autoradiogram (top) and Coomassie-staining (bottom) are as described in **A**.

Since purified tubulin is composed of α-and β-tubulin heterodimers, we sought to determine which species is methylated by SET8. Recombinant fusion proteins of either α-tubulin or β-tubulin with MBP were purified and incubated with SET8 along with the radioactive methyl donor. Upon incubation of SET8 with α/β tubulin and AdoMet, both SET8 and tubulin(s) were labeled. However, only MBP-α-tubulin, but not MBP-β-tubulin was methylated, along with SET8 itself, when the individual recombinant proteins were tested (Fig 2B). These data indicate that α-tubulin is the target for SET8. Mass spectrometry was used to determine which lysine residue(s) of α-tubulin were methylated by SET8. In the control samples without exogenous SET8, lysine methylation of α-tubulin on K304, and of β-tubulin on K19 and K297 were detected (Appendix Fig S2A), none of which have previously been reported. As anticipated from the previous data (Fig 2B), incubation with exogenous SET8 did not induce detectable methylation at any other sites on β-tubulin. However, SET8 did induce methylation of three additional lysine residues of α-tubulin - K280, K311 and K352 - which were all monomethylated (Fig 2C, Appendix Fig S2A). Of these three lysines, only K311 is located on the outside surface of MTs, whereas K352 is at the interface between α-tubulin and the β-tubulin in the adjacent heterodimer, and K280 is on the inside surface of MTs (Fig 2D, Appendix Fig S2B,C). In addition, only the sequence surrounding K311 (RHGK_311_) resembles those of other known SET8 target sequences: histone H4 (RHRK_20_) and p53 (RHKK_382_) (Shi *et al.*, 2007). In contrast, the sequences of the other α-tubulin sites, SAEK_280_ and TGFK_352_, do not resemble other known physiological SET8 tarets. To determine the relative efficiency of targeting these lysines *in vitro*, each was independently mutated in the context of the full-length MBP-α-tubulin, and purified proteins were tested for incorporation of radioactivity upon incubation with SET8. In order to test the relevant degree of methylation by SET8 at the identified sites, each lysine was mutated indivually to serine, maintaining a similar structure and hydrophilicity, but removing the charge. Consistent with K311 being the best sequence match with other SET8 targets, only mutation of K311 was no longer detectably modified by SET8, whereas mutation of K280, K304, or K352 did not appreciably affect the amount of methylation of the substrates (Fig 2E).

In order to test further the inherent targeting of the various α-tubulin sites by SET8, peptides spanning these three sites (K280, K311, K352), as well as K40, reported to be methylated by SETD2 (Park *et al.*, 2016), and K304, modified in purified porcine tubulin (Appendix Fig S2A), were incubated with purified wild type SET8 *in vitro*. Only the K311-containing peptide was robustly methylated (Appendix Fig S3A, Appendix Table S2). In addition, radioactive incorporation into the K311-containing peptide was absent when incubated with catalytically inactive SET8 (D338A) *in vitro*, and methylation abolished if the K311 residue was either mutated (K311A, K311S) or already modified (K311Me, K311Ac) (Appendix Fig S3B, Appendix Table S2). Although the *in vitro* targeting of the α-tubulin K311-containing peptide by SET8 is robust (Appendix Fig S3C), SET8 methylates histone H4 much more efficiently, consistent with the ability of Histone H4 to strongly compete against tubulin for methylation by SET8 (Fig 2A). In contrast, but consistent with the previous report (Park *et al.*, 2016), methylation by the only other reported tubulin methyltransferase, SETD2, of the α-tubulin K40-containing peptide was not detectable over background, despite some methylation of histone H3 by this enzyme *in vitro* (Appendix Fig S3D).

Taken together, these observations indicated that SET8 methyltransferase has the capacity to directly methylate α-tubulin, particularly at K311.

### Transcription factor LSF associates with both SET8 and tubulin

DNA-binding proteins recruit chromatin writers to modify histones (Brownell *et al.*, 1996;Hassig and Schreiber, 1997;Struhl, 1999;Taunton *et al.*, 1996), suggesting the possibility that tubulin-binding proteins might similarly recruit SET8 to target sites on microtubules resulting in tubulin modification. Our previous studies showed that the transcription factor LSF interacts with DNMT1, and addition of an inhibitor of the LSF-DNMT1 interaction resulted in alterations in the genomic DNA methylation profile (Chin *et al.*, 2016); this is consistent with recruitment of DNMT1 to DNA by LSF in order to facilitate DNA methylation at specific sites. Since DNMT1 complexes with SET8, and both SET8 and LSF (Rajasekaran *et al.*, 2015) are required for mitotic progression, we proposed the novel hypothesis that the transcription factor LSF might also recruit SET8 to microtubules in order to facilitate α-tubulin methylation by SET8. In support of this hypothesis, there is precedent for some DNA-binding transcription factors also binding microtubules (Alexandrova *et al.*, 1995;Dong *et al.*, 2000;Giannakakou *et al.*, 2000;Maxwell *et al.*, 1991;Niklinski *et al.*, 2000;Ziegelbauer *et al.*, 2001), although for the purpose of sequestering the transcription factors in the cytoplasm and/or facilitating their transport into the nucleus.

To test this hypothesis, multiple assays were initially performed to evaluate whether LSF interacts with SET8 and tubulin(s). *In vitro*, direct interaction between recombinant, purified SET8 and purified LSF was evaluated by a GST pull-down assay (Fig 3A). Using fusion proteins between GST and either full length SET8, or its N- or C-terminal fragments, the His-LSF bound specifically to the N-terminal region of SET8 (Fig 3A), the same domain that bound purified tubulin (Fig 1F). The binding of purified α/β-tubulin to purified GST-LSF was also evaluated and mapped to specific regions within LSF. Both α- and β-tubulins showed similar binding profiles to the panel of LSF fusion proteins, as expected given their stable heterodimeric structure (Fig 3B). Binding of tubulins to the full-length GST-LSF was greater than to the control GST, although weak compared to some of the other fusion proteins, due to the sensitivity of the full-length GST-LSF fusion protein to cleavage in bacterial culture, resulting in significant purification of the GST domain alone in the preparation. However, the tubulins interacted strongly with two specific domains of LSF whose GST fusion proteins were stable: the DNA binding domain (DBD), and to a lesser extent, the SAM domain (Fig 3B). Further analysis suggests that it is the C-terminal portion of the DBD that contains the tubulin interaction surface in this domain, since the GST-LSF 2 protein also binds both tubulins to a high degree. Finally, purified His-LSF also interacted in parallel assays with purified recombinant full-length GST-α-tubulin (Fig 3C). These *in vitro* protein-protein interaction results indicate that all pairwise interactions among SET8, LSF, and α-tubulin occur through direct binding with each other.

**Figure 3.**
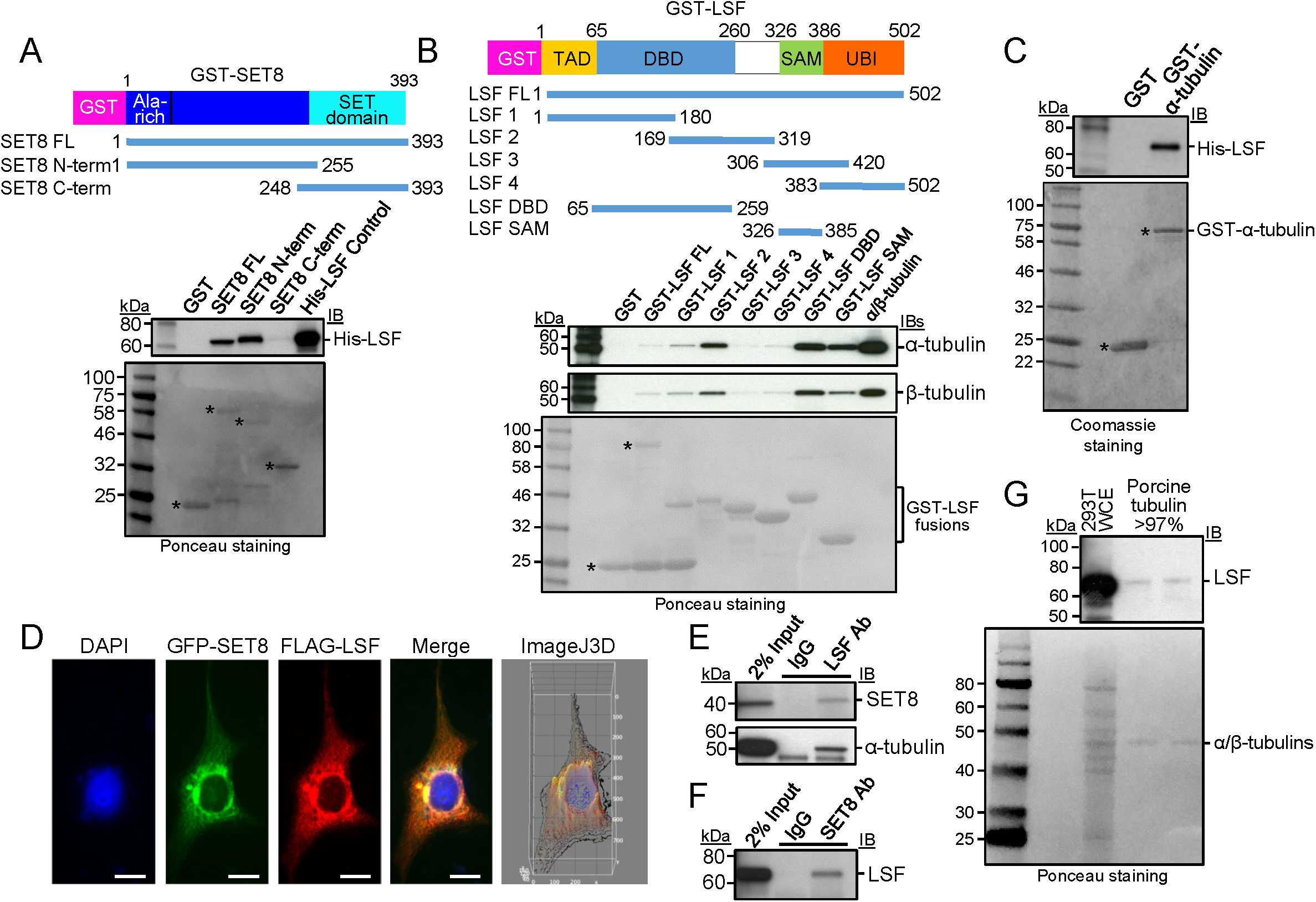
LSF interacts directly with SET8 and tubulin. **A** GST-pull down analysis of purified His-LSF to full-length or the indicated overlapping segments of SET8 fused to GST. Top: immunoblot (IB) in which biotinylated molecular weight markers are visualized; bottom: Ponceau staining of the same gels (shown in grayscale) in which standard molecular weight markers are visualized. *Expected positions of migration of the GST-proteins. **B** GST-pull down analysis of purified porcine tubulin to purified full-length or the indicated overlapping segments of LSF fused to GST. Gels are as described in **A**. * or bracket: Expected positions of migration of the GST-proteins. **C** GST-pull down analysis of recombinant, purified His-tagged LSF to purified α-tubulin fused to GST. Gels are as described in **A**, except that the protein gel was stained with Coomassie. **D** Plasmids expressing 3xFLAG-LSF and GFP-SET8 were transfected into COS7 cells. Anti-FLAG antibody was visualized with a red fluorescing secondary antibody, and DNA was visualized with DAPI. The merged image indicates colocalization of GFP-SET8 with FLAG-LSF (yellow), concentrated largely near the nuclear membrane (Manders correlation coefficient of LSF and SET8 colocalization is 0.9, as determined via the Image J 3D analysis). The majority of overexpressed 3XFlagLSF was cytoplasmic with only a minority detected in the nucleus. **E** Specific co-immunoprecipitation of endogenous SET8 (top) and endogenous α-tubulin (bottom) from HEK293 cellular extracts, using antibodies to LSF as compared to control IgG. **F** Specific co-immunoprecipitation of endogenous LSF from HEK293 cellular extracts, using antibodies to SET8 as compared to control IgG. **G** Immunoblotting of purified porcine brain tubulin (rPeptide, >97%) shows the presence of LSF, using a LSF monoclonal antibody. Representative also of results obtained using a separate source of purified tubulin: MP Biomedical >99%. Positive control for LSF migration: 293T WCE (HEK293T cell whole cell extract). Top: immunoblot; bottom: Ponceau staining using standard molecular weight markers.

To examine whether interactions of LSF with both SET8 and tubulin also take place in cells, multiple approaches were taken. First, upon co-expression of GFP-SET8 and 3xFlag-tagged LSF in transient transfection assays, the two proteins significantly co-localized, predominantly in the cytoplasm (Fig 3D). Second, the presence of complexes between endogenous cellular proteins were tested by co-immunoprecipitation experiments using lysates from the human HEK293 cell line. With antibodies against endogenous LSF, but not control antibodies, both endogenous SET8 and endogenous α-tubulin co-immunoprecipitated with LSF (Fig 3E). Reciprocally, SET8 antibodies, but not control antibodies, also co-immunoprecipitated endogenous LSF (Fig 3F). Finally, the possibility of relevant LSF-tubulin interactions was investigated by analyzing whether LSF was present in commercial, highly purified tubulin preparations. These preparations are obtained in part by multiple rounds of polymerization/depolymerization of the tubulin, and are more than 97-99% pure. They are well known to contain additional proteins that are defined as microtubule-associated proteins (MAPs). Immunoblots using a LSF monoclonal antibody did indeed detect a band comigrating with LSF, albeit at a very low level (Fig 3G). LSF was reproducibly detected in this manner in multiple commercially purified preparations of tubulin.

Taken together, these results demonstrate that LSF interacts directly with both SET8 and α-tubulin *in vitro*, and also associates with both *in vivo*. Furthermore, LSF, although a transcription factor, appears to be a previously unidentified MAP.

### LSF promotes tubulin methylation by SET8

The demonstration of pairwise, physical interactions between LSF, tubulin, and SET8 set the stage for directly testing the hypothesis that LSF could mediate the methylation of α-tubulin by SET8. Thus, recombinant GST-SET8 and the methyl donor were incubated with tubulin in the presence of increasing concentrations of purified His-LSF (Fig 4A). Tubulin methylation increased upon increasing LSF from a 1:4 to 2:1 molar ratio of LSF:GST-SET8, suggesting that LSF can mediate tubulin methylation by SET8. Note that in this experiment, there was more SET8 relative to tubulin than in other experiments (Fig 4A, bottom), resulting in a greater degree of automethylation of SET8 compared to tubulin methylation (Fig 4A, top), presumably due to substrate competition between SET3 itself and α-tubulin. A similar experiment was performed using recombinant MBP-α-tubulin as substrate for SET8, which also showed that increasing levels of LSF enhanced methylation of the MBP-α-tubulin (Appendix Fig S4A).

**Figure 4.**
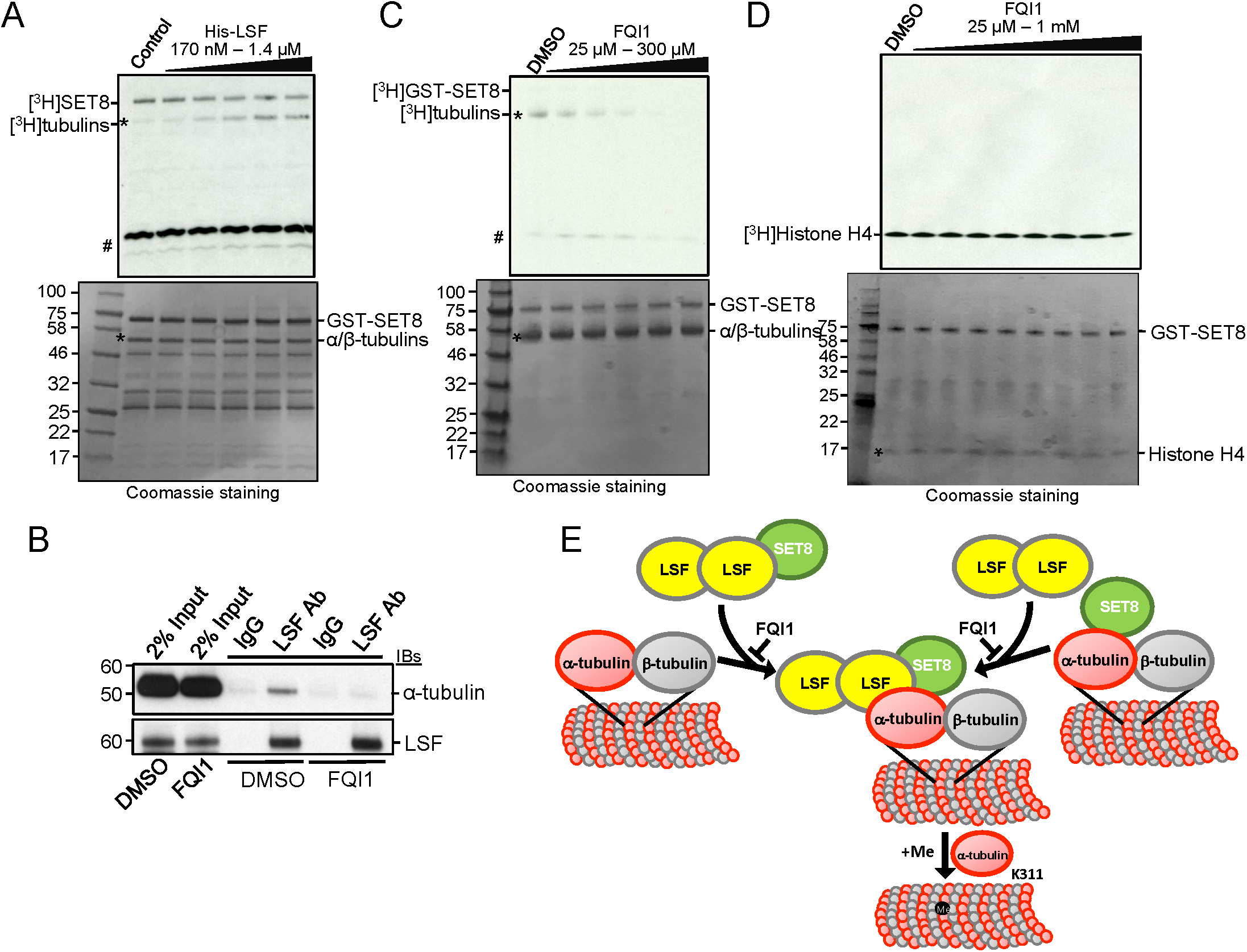
LSF and FQI1 oppositely affect methylation of tubulin by SET8. **A** Tubulin (>99%, MP Biomedical) methylation reactions were performed with addition of the indicated, increasing range of concentrations of LSF. Top: autoradiogram of methyltransferase assays, showing methylation of tubulin (*) and automethylation of GST-SET8. The higher relative levels of GST-SET8 to α/β-tubulin in this experiment led to greater initial automethylation of SET8 relative to tubulin methylation. # indicates the migration of [^3^H]-labeled impurities. Bottom: Coomassie-staining of the same gel (shown in grayscale) indicating relative levels of the components in the reaction. As in Fig 2B, protein bands <50 kDa are from the purified GST-SET8 preparation, and are more evident in this experiment. **B** Co-immunoprecipitation of endogenous α-tubulin with endogenous LSF from HEK293T cell lysates was disrupted upon treatment of the cells with 2.5 μM FQI1 for 24 hr. **C** Tubulin (>99%, MP Biomedical) methylation reactions were performed with addition of the indicated, increasing range of concentrations of FQI1. At 100 µM FQI1 (lane 4), methylation is decreased ~3-fold. Gels are labeled as in **A**. **D** Histone H4 methylation reactions at limiting amounts of histone H4 (200 ng) were performed with addition of the indicated, increasing range of concentrations of FQI1. Gels are labeled as in **A**. **E.** Model for the recruitment and/or activation of SET8 at microtubules by LSF, and the subsequent methylation of α-tubulin by SET8.

The LSF small molecule inhibitor, FQI1, inhibits LSF binding to DNA (Grant *et al.*, 2012), as well as binding of LSF to certain protein partners (Chin *et al.*, 2016). To determine whether FQI1 would also diminish the interaction in cells between LSF and tubulin, cell lysates from vehicle-versus FQI1-treated cells were analyzed by co-immunoprecipitation assays. These demonstrated a significant reduction in the LSF-tubulin interaction after FQI1 incubation (Fig 5B). Since FQI1 can inhibit the LSF-tubulin interaction *in vivo*, it was used to interrogate whether the interaction of LSF and tubulin was important for stimulating the SET8-mediated methylation of tubulin *in vitro*. Since LSF is already present in the tubulin preparations, initially, FQI1 was added to reactions containing only SET8, methyl donor, and purified tubulin. Tubulin methylation decreased with increasing concentrations of FQI1 (Fig 5C), consistent with the presence of LSF and its ability to enhance SET8-dependent tubulin methylation. Whether FQI1 specifically inhibits LSF in these assays was tested in two ways. First, it was demonstrated that the presence of FQI1 prevented any increase in tubulin methylation upon addition of purified His-LSF (Appendix Fig S4B, compare lanes 7 and 8). Second, the possibility that FQI1 directly inhibits SET8 catalytic activity was tested using Histone H4, instead of α-tubulin, as a substrate. Limiting amounts of histone H4 were added in this experiment to enhance the sensitivity of the assay. FQI1 did not inhibit methylation of histone H4 by SET8, in contrast to its effect on α-tubulin methylation (Fig 5D). In addition, when SET8 was incubated with whole cell extract in the presence of the radioactive methyl donor, FQI1 did not appreciably diminish methylation of any of the other proteins in the extract (Appendix Fig S4B, compare lanes 1 and 2).

These data indicate that LSF enhances tubulin methylation by SET8, and conversely that FQI1, which abrogates the tubulin-LSF interaction (Fig 4B), impedes methylation of α-tubulin by SET8. Overall, these data support a model that LSF recruits SET8 to tubulin, and/or that LSF binding as a ternary complex to SET8 and tubulin activates the methylase activity of SET8 already associated with tubulin (Fig 4E). To probe whether LSF recruits SET8 to tubulin, we again used FQI1 to disrupt the LSF-tubulin interactions. FQI1 did diminish co-immunoprecipitation of endogenous tubulin with SET8 antibodies (Appendix Fig S4C), supporting the recruitment model. However, a caveat to this straightforward interpretation was that SET8 immunoprecipitation was also somewhat diminished, although less so, after incubation of the cells with FQI1.

## Discussion

A number of post-translational modifications of microtubules are well established, including the enzymes responsible for these modifications. However, thus far, only limited insights have been obtained regarding their biological roles, and tubulin PTMs have generally remained less amenable to straightforward *in vitro* functional studies. A number of posttranslational modifications of tubulins have been reported, along with their modifying enzymes, including acetylation of lysine 40 in α-tubulin by αTAT1 (MEC-17 in *C. elegans*), deacetylation of the same residue by Sirt2 (Sirtuin type2), HDAC5, and HDAC6, and polyglutamylation of both α- and β-tubulins at multiple C-terminal glutamate residues by TTLL4, 5, and 7 (Ikegami *et al.*, 2006;Janke *et al.*, 2005). Although extensive research has been carried out on various modified sites and has identified the relevant enzymes, only one study has previously identified lysine methylation of tubulin (Park *et al.*, 2016), which is the focus of this report. Walker’s group reported that SETD2, known as a histone methyltransferase for a chromatin activation mark, H3K36me3, also methylates α-tubulin at K40. Furthermore, the loss of SETD2-mediated tubulin methylation resulted in mitotic microtubule defects and genomic instability. Here, we describe a distinct, novel lysine methylation of α-tubulin at K311 and identify the enzyme responsible for its modification as SET8. In addition, we identify methylation of β-tubulin purified from mammalian brain at K19 and K297, and are pursuing identification of the enzymes responsible.

Importantly, we also demonstrate the surprising finding that a transcription factor, LSF, moonlights as a microtubule-associated protein, and that it can recruit SET8 to tubulin and/or enhance its activity on tubulin to facilitate the modification. The recruitment mechanism mirrors mechanisms of targeting of histone writers to chromatin, expanding the model of the parallel nature between the generation of the histone and tubulin codes. Furthermore, these data indicate that transcription factors more generally may be able to regulate tubulin modifications, and thereby microtubule dynamics. Although several transcription factors have previously been reported to bind microtubules, including c-myc (Alexandrova *et al.*, 1995;Niklinski *et al.*, 2000), MIZ-1 (Ziegelbauer *et al.*, 2001), p53 (Giannakakou *et al.*, 2000;Maxwell *et al.*, 1991), and Smads (Dong *et al.*, 2000), in all these cases the biological relevance proposed or demonstrated was to sequester the transcription factors in the cytoplasm, and/or to help transport the transcription factor into the nucleus. Thus, all previous indications had to do with regulation of transcription activity, not of microtubule function. This is the first case in which binding of a transcription factor directly to microtubules would lead to altered microtubule modifications, and to altered function. We propose that this presents a new paradigm that may well occur with other transcription factors, as well.

Tubulin PTMs are generally thought to regulate protein-protein interactions within the microtubule cytoskeleton, thereby regulating signaling events in the cell. To date, a large variety of microtubule associated proteins (MAPs) have been characterized, many of which stabilize and destabilize microtubules, are associated with the coupling of molecular motors and microtubules, and play critical roles in spindle formation (Stanton *et al.*, 2011).

Proper mammalian cell cycle progression requires precise modulation of SET8 levels, which suggested that SET8 and H4K20me1 function as novel regulators of cell cycle progression, although the previous focus has been on regulation of S phase (Milite *et al.*, 2016). With our demonstration that SET8 can also methylate α-tubulin, the roles of non-histone substrates must also be considered as causes for SET8-mediated regulation of the cell cycle, and in particular of mitosis when SET8 is most abundant (Rice *et al.*, 2002). As in mammals, SET8 is essential for life in *D. melanogaster*, as SET8 mutants die during larval development. However, upon replacement of all *Drosophila* histone H4 genes with multiple copies of a mutant histone H4K20A, the flies survived to adulthood without apparent DNA replication defects (McKay *et al.*, 2015). Thus, contrary to the prevailing view, histone H4 was not the critical target for SET8. Given the minimal biological effects in *Drosophila* of mutating histone H4K20, we propose that α-tubulin methylation is a strong candidate for mediating critical SET8 consequences. Notably, *D. melanogaster* and human α-tubulins are 98% identical with all the lysines throughout the sequence being conserved.

Mitosis is viewed as a vulnerable target for inhibition in cancer (Komlodi-Pasztor *et al.*, 2012). In that light, it is notable that LSF promotes oncogenesis in hepatocellular carcinoma, the sixth most common cancer worldwide and the second highest cause of cancer-related death globally (Laursen, 2014). LSF is overexpressed in human hepatocellular carcinoma cell lines, and over 90% of human hepatocellular carcinoma patient samples, showing significant correlation with stages and grades of the disease (Yoo *et al.*, 2010). The lead LSF inhibitor, FQI1, induces apoptosis in an aggressive hepatocellular carcinoma cell line *in vitro* and significantly inhibits tumor growth in multiple mouse hepatocellular carcinoma models, with no observable toxicity to normal tissues (Grant *et al.*, 2012;Rajasekaran *et al.*, 2015). These new findings that LSF interacts with tubulin and SET8, and that FQI1 disrupts the LSF-tubulin interaction, may be related to the impact of the LSF inhibitors in hepatocellular carcinoma cells and tumors. Expression both of particular tubulins (e.g. TUBA1B) and of SET8 are upregulated in hepatocellular carcinoma tumor samples, as compared to normal liver (Guo *et al.*, 2012;Lu *et al.*, 2013). Moreover, SET8 is required to maintain the malignant phenotype of various cancer types (Hou *et al.*, 2016). Given the current lack of effective treatments, further investigation into the relevance of the LSF-tubulin-SET8 pathway to hepatocellular carcinoma and other cancer types in which LSF is oncogenic may aid in targeted and effective treatment.

## Material and Methods

### Cell Culture, immunoprecipitation, and immunofluorescence

HEK293T and COS7 cells were cultured in DMEM media supplemented with 10% fetal bovine serum. FQI1 (Millipore/Sigma #438210) treatment of HEK293T cells was for 24 hours at 37°C with 2.5 μM FQI1.

Immunoprecipitation (IP) and immunofluorescence experiments were carried out as described previously (Andrews and Faller, 1991;Estève *et al.*, 2006). For the immunoprecipitation, 1 mg of total HEK293T cellular extract was incubated with 5 μg of anti-SET8 antibody (Active Motif, 61009), anti-LSF antibody (Millipore,17-10252), or mouse anti-FLAG antibody (F1804, Sigma-Aldrich). The immunoprecipitates were blotted with anti-α-tubulin, anti-β-tubulin (Sigma T9026, T8328), anti-SET8 (Abcam, ab3798), anti-Pr-Set7 (D11) (Santa Cruz, sc-377034), anti-LSF (BD Biosciences) antibodies or anti-FLAG (CST, 14793), as per the manufacturer’s dilution recommendations. Cellular extracts were also immunoprecipitated with normal IgG (Cell Signaling Technology) as a negative control for all IP experiments.

For immunofluorescence to detect α-tubulin and SET8 co-localization, COS7 cells were grown on coverslips and transfected with a GFP-SET8 expression plasmid. After cells were fixed with paraformaldehyde, the cells were incubated with anti-α-tubulin and the microtubules visualized with an anti-mouse IgG coupled with Alexa Fluor 488 (Molecular Probes) using a confocal microscope (Zeiss LSM510). For the detection of SET8 and LSF co-localization, COS7 cells were co-transfected with GFP-SET8 and 3XFlag-LSF expression plasmids; the epitope tagged LSF was detected by mouse anti-FLAG antibody (F3165, Sigma-Aldrich) and visualized with an anti-mouse IgG coupled with Alexa Fluor 488 (Molecular Probes). DAPI was used to stain nuclear DNA. For analysis of FQI1 effects on mitosis, paraformaldehyde-fixed cells were analyzed for microtubules with anti-α-tubulin antibody (Abcam #AB7750) and stained with DAPI (Invitrogen) for visualizing genomic DNA. Samples were imaged using a Zeiss Axioimager M1 microscope utilizing 63× and 100× magnifications.

### GST and MBP pull down assays

LSF, SET8 and α-tubulin cDNAs were cloned into the pGEX-5X-1 (GE Healthcare) or pMalC4X (New England Biolabs) vector and GST-tagged or MBP-tagged proteins were captured using Glutathione Sepharose beads (GE Healthcare) or amylose resin (New England Biolabs), respectively. Sepharose beads containing approximately 10 μg of fusion protein were incubated for 2 hours at 4°C with purified tubulin (MP-biomedical), recombinant His-tagged LSF, or recombinant His-tagged SET8, the latter two being purified from *E. coli*. Proteins bound to the beads were resolved by 10-20% SDS-PAGE. LSF, SET8 and α-tubulin were visualized by immunoblotting by using anti-LSF (BD Biosciences), anti-SET8 (Active Motif) or anti-α-tubulin (Sigma-Aldrich), respectively.

### *In vitro* methylation assays

Approximately 1 μg of recombinant GST-SET8 (in 50% glycerol) and 2 μg of the purified tubulin (MP-Bioscience, #0877115), recombinant MBP-α-tubulin, or recombinant MBP-β-tubulin were incubated with 6 μM radioactively labeled [^3^H] AdoMet (Perkin Elmer #NET155V001MC) in 1X HMT buffer containing 5 mM Tris pH 8.0, 0.5 mM DTT at room temperature for overnight. As indicated, histone H4 (NEB #2504S), recombinant His-LSF protein, or FQI1 inhibitor were added to the reaction. Samples were separated by electrophoresis through a 10% Tricine Gel (Invitrogen) and the gel was stained with Coomassie Brilliant Blue (shown in grayscale in the figures) and incubated with EN3HANCE (PerkinElmer) solution. The gel was dried and exposed to autoradiography film for 1 week. For the peptide assays, the specific peptides of α-tubulin were synthesized from AnaSpec Inc. Sequences are listed in Appendix Table S2. Two μg of each peptide and 2 μg of purified wild type or mutant SET8 or full-length SETD2 (Active Motif) were incubated with radioactively labeled [^3^H] AdoMet at room temperature overnight. Samples were spotted onto P81 filters (Whatman 3698325) and the filters were washed 3 times with 0.3 M ammonium bicarbonate. The level of incorporated [^3^H]CH_3_ was determined using liquid scintillation counting.

### Mass-spectrometric analysis

For identification of proteins in the tubulin preparations, >97% purified porcine tubulin (rPeptide, # T-1201-1) and >99% purified bovine tubulin (MP-Bioscience, #0877112) were separated by electrophoresis through a 10-20% Tris-Glycine polyacrylamide gel. The gel was stained with Coomassie Blue; sections of the gels around 55 kDa, and below 55 kDa, respectively, were excised (Appendix Fig S1A). Excised gel band were analyzed by the Taplin Mass Spectrometry Facility, Harvard Medical School. For identification of tubulin modifications, purified tubulin (MP-Bio, #0877115) was incubated with nonradioactive AdoMet, with or without recombinant GST-SET8, overnight at room temperature and the samples were separated by electrophoresis through a 10% Tris-glycine gel. Excised gel bands were digested with either subtilisin or trypsin in 0.01% ProteaseMax in 50 mM NH4HCO3 for 1 hr at 50°C. Digestion was quenched with trifluoroacetic acid and samples were dried. Each digest was individually reconstituted and analyzed by direct injection onto an analytical column 25 cm 100 µm ID Aqua 3 µm with Easy n1000 nLC-QExactive at 300 nL/min. Acquired HCD spectra were searched using Proteome Discoverer 2.0 and SWISSPROT June 2015 database using nonspecific cleavage and Cys = 57.02146 static modification. Dynamic modifications were set for Met = 15.99492. Searches were semi-specific with K= 14.016 dynamic modification. Results were filtered with Percolator for high confidence spectrum matches.

### Biochemical fractionation of HEK293T cells

Subcellular fractions from cultured HEK293T cells were obtained using the Cell Fractionation Kit (CST, #9038) according to the manufacturer’s protocol. Cytoplasm, nuclear, and whole cell extracts were separated by electrophoresis through a 10-20% 10-20% Tris-Glycine and the resulting membrane immunoblotted with anti-MEK1/2 (D1A5, #8727), Anti-histone H3(CST, #9715), anti-β-actin (CST, #4970), anti-SET8 (Active Motif, #61009).

## Acknowledgements

We thank W. Jack, C. Carlow, and G.M. Cooper for critical reading of the manuscript, D. Comb, Sir R.J. Roberts and J. Ellard for encouragement, and L. Brown for the preparation of FQI1. Basic research support for H.G.C., P.O.E, C.R. and S.P. was provided by New England Biolabs, Inc. U.H. was supported by Ignition Awards from Boston University and a Johnson & Johnson Clinical Innovation Award through Boston University. S.E.S. was supported by the NIH (R01 GM078240 and R24 GM111625).

## Author Contributions

H.G.C. conceived the initial idea and designed and performed most of the experiments. P.O.E. performed the microscopy and imaging. C.R. performed the mass spectrometry analysis. J.L. contributed to discussions and suggested guidelines for mass spectroscopy. S.E.S. provided information on storage and usage of FQI1. U.H. and S.P. supervised the work. H.G.C., U.H., and S.P. wrote the manuscript, which was reviewed by all authors.

## Conflict of Interest

The authors declare no conflicts of interest.

## References

Alexandrova N, Niklinski J, Bliskovsky V, Otterson GA, Blake M, Kaye FJ, Zajac-Kaye M (1995) The N-terminal domain of c-Myc associates with α-tubulin and microtubules in vivo and in vitro. Mol Cell Biol, 15, 5188–5195.

Andrews NC, Faller DV (1991) A rapid micropreparation technique for extraction of DNA-binding proteins from limiting numbers of mammalian cells. Nucleic Acids Res, 19, 2499.

Brownell JE, Zhou J, Ranalli T, Kobayashi R, Edmondson DG, Roth SY, Allis CD (1996) Tetrahymena histone acetyltransferase A: A homolog to yeast Gcn5p linking histone acetylation to gene activation. Cell, 84, 843–851.

Chin HG, Ponnaluri VKC, Zhang G, Estève P-O, Schaus SE, Hansen U, Pradhan S (2016) Transcription factor LSF-DNMT1 complex dissociation by FQI1 leads to aberrant DNA methylation and gene expression. Oncotarget, 7, 83627–83640.

Dillon SC, Zhang X, Trievel RC, Cheng X (2005) The SET-domain protein superfamily: protein lysine methyltransferases. Genome Biol, 6, 227.

Dong C, Li Z, Alvarez R, Jr., Feng X-H, Goldschmidt-Clermont PJ (2000) Microtubule binding to Smads may regulate TGFβ activity. Mol Cell, 5, 27–34.

Estève P-O, Chin HG, Smallwood A, Feehery GR, Gangisetty O, Karpf AR, Carey MF, Pradhan S (2006) Direct interaction between DNMT1 and G9a coordinates DNA and histone methylation during replication. Genes Dev, 20, 3089–3103.

Fan R-H, Li J, Wu N, Chen P-S (2011) Late SV40 factor: A key mediator of Notch signaling in human hepatocarcinogenesis. World J Gastroenterol, 17, 3420–3430.

Fang J, Feng Q, Ketel CS, Wang H, Cao R, Xia L, Erdjument-Bromage H, Tempst P, Simon JA, Zhang Y (2002) Purification and functional characterization of SET8, a nucleosomal histone H4-lysine 20-specific methyltransferase. Curr Biol, 12, 1086–1099.

Giannakakou P, Sackett DL, Ward Y, Webster KR, Blagosklonny MV, Fojo T (2000) p53 is associated with cellular microtubules and is transported to the nucleus by dynein. Nat Cell Biol, 2, 709–717.

Grant TJ, Bishop JA, Christadore LM, Barot G, Chin HG, Woodson S, Kavouris J, Siddiq A, Gredler R, Shen X-N, Sherman J, Meehan T, Fitzgerald K, Pradhan S, Briggs LA, Andrews WH, Sarkar D, Schaus SE, Hansen U (2012) Antiproliferative small molecule inhibitors of transcription factor LSF reveal oncogene addiction to LSF in hepatocellular carcinoma. Proc Natl Acad Sci USA, 109, 4503–4508.

Gu Y, Li H, Zhao L, Zhao S, He W, Rui L, Su C, Zheng H, Su R (2015) GRP78 confers the resistance to 5-FU by activating the c-Src/LSF/TS axis in hepatocellular carcinoma. Oncotarget, 6, 33658–33674.

Guo Z, Wu C, Wang X, Wang C, Zhang R, Shan B (2012) A polymorphism at the miR-502 binding site in the 3’-untranslated region of the histone methyltransferase *SET8* is associated with hepatocellular carcinoma outcome. Int J Cancer, 131, 1318–1322.

Hansen U, Owens L, Saxena UH (2009) Transcription factors LSF and E2Fs: Tandem cyclists driving G0 to S? Cell Cycle, 8, 2146–2151.

Hassig CA, Schreiber SL (1997) Nuclear histone acetylases and deacetylases and transcriptional regulation: HATs off to HDACs. Curr Opin Chem Biol, 1, 300–308.

Hou L, Li Q, Yu Y, Li M, Zhang D (2016) SET8 induces epithelial-mesenchymal transition and enhances prostate cancer cell metastasis by cooperating with ZEB1. Mol Med Rep, 13, 1681–1688.

Ikegami K, Mukai M, Tsuchida J, Heier RL, Macgregor GR, Setou M (2006) TTLL7 is a mammalian β-tubulin polyglutamylase required for growth of MAP2-positive neurites. J Biol Chem, 281, 30707–30716.

Janke C (2014) The tubulin code: molecular components, readout mechanisms, and functions. J Cell Biol, 206, 461–472.

Janke C, Rogowski K, Wloga D, Regnard C, Kajava AV, Strub J-M, Temurak N, van Dijk J, Boucher D, van Dorsselaer A, Suryavanshi S, Gaertig J, Eddé B (2005) Tubulin polyglutamylase enzymes are members of the TTL domain protein family. Science, 308, 1758–1762.

Kim JS, Son SH, Kim MY, Choi D, Jang I-S, Paik SS, Chae JH, Uversky VN, Kim CG (2017) Diagnostic and prognostic relevance of CP2c and YY1 expression in hepatocellular carcinoma. Oncotarget, 8, 24389–24400.

Komlodi-Pasztor E, Sackett DL, Fojo AT (2012) Inhibitors targeting mitosis: tales of how great drugs against a promising target were brought down by a flawed rationale. Clin Cancer Res, 18, 51–63.

Kotarba G, Krzywinska E, Grabowska AI, Taracha A, Wilanowski T (2018) TFCP2/TFCP2L1/UBP1 transcription factors in cancer. Cancer Lett, 420, 72–79.

Laursen L (2014) A preventable cancer. Nature, 516, S2–S3.

Lu C, Zhang J, He S, Wan C, Shan A, Wang Y, Yu L, Liu G, Chen K, Shi J, Zhang Y, Ni R (2013) Increased α-tubulin1b expression indicates poor prognosis and resistance to chemotherapy in hepatocellular carcinoma. Dig Dis Sci, 58, 2713–2720.

Maxwell SA, Ames SK, Sawai ET, Decker GL, Cook RG, Butel JS (1991) Simian virus 40 large T antigen and p53 are microtubule-associated proteins in transformed cells. Cell Growth Differ, 2, 115–127.

McKay DJ, Klusza S, Penke TJR, Meers MP, Curry KP, McDaniel SL, Malek PY, Cooper SW, Tatomer DC, Lieb JD, Strahl BD, Duronio RJ, Matera AG (2015) Interrogating the function of metazoan histones using engineered gene clusters. Dev Cell, 32, 373–386.

Milite C, Feoli A, Viviano M, Rescigno D, Cianciulli A, Balzano AL, Mai A, Castellano S, Sbardella G (2016) The emerging role of lysine methyltransferase SETD8 in human diseases. Clin Epigenetics, 8, 102.

Niklinski J, Claassen G, Meyers C, Gregory MA, Allegra CJ, Kaye FJ, Hann SR, Zajac-Kaye M (2000) Disruption of Myc-tubulin interaction by hyperphosphorylation of c-Myc during mitosis or by constitutive hyperphosphorylation of mutant c-Myc in Burkitt’s lymphoma. Mol Cell Biol, 20, 5276–5284.

Nishioka K, Rice JC, Sarma K, Erdjument-Bromage H, Werner J, Wang Y, Chuikov S, Valenzuela P, Tempst P, Steward R, Lis JT, Allis CD, Reinberg D (2002) PR-Set7 is a nucleosome-specific methyltransferase that modifies lysine 20 of histone H4 and is associated with silent chromatin. Mol Cell, 9, 1201–1213.

Park IY, Powell RT, Tripathi DN, Dere R, Ho TH, Blasius TL, Chiang YC, Davis IJ, Fahey CC, Hacker KE, Verhey KJ, Bedford MT, Jonasch E, Rathmell WK, Walker CL (2016) Dual Chromatin and Cytoskeletal Remodeling by SETD2. Cell, 166, 950–962.

Powell CMH, Rudge TL, Zhu Q, Johnson LF, Hansen U (2000) Inhibition of the mammalian transcription factor LSF induces S-phase-dependent apoptosis by downregulating thymidylate synthase expression. EMBO J, 19, 4665–4675.

Rajasekaran D, Siddiq A, Willoughby JLS, Biagi JM, Christadore LM, Yunes SA, Gredler R, Jariwala N, Robertson CL, Akiel MA, Shen X-N, Subler MA, Windle JJ, Schaus SE, Fisher PB, Hansen U, Sarkar D (2015) Small molecule inhibitors of Late SV40 Factor (LSF) abrogate hepatocellular carcinoma (HCC): Evaluation using an endogenous HCC model. Oncotarget, 6, 26266–26277.

Rice JC, Nishioka K, Sarma K, Steward R, Reinberg D, Allis CD (2002) Mitotic-specific methylation of histone H4 Lys 20 follows increased PR-Set7 expression and its localization to mitotic chromosomes. Genes Dev, 16, 2225–2230.

Seol HS, Lee SE, Song JS, Rhee J-K, Singh SR, Chang S, Jang SJ (2016) Complement proteins C7 and CFH control the stemness of liver cancer cells via LSF-1. Cancer Lett, 372, 24–35.

Shi X, Kachirskaia I, Yamaguchi H, West LE, Wen H, Wang EW, Dutta S, Appella E, Gozani O (2007) Modulation of p53 function by SET8-mediated methylation at lysine 382. Mol Cell, 27, 636–646.

Song Y, Brady ST (2015) Post-translational modifications of tubulin: pathways to functional diversity of microtubules. Trends Cell Biol, 25, 125–136.

Stanton RA, Gernert KM, Nettles JH, Aneja R (2011) Drugs that target dynamic microtubules: a new molecular perspective. Med Res Rev, 31, 443–481.

Struhl K (1999) Fundamentally different logic of gene regulation in eukaryotes and prokaryotes. Cell, 98, 1–4.

Taunton J, Hassig CA, Schreiber SL (1996) A mammalian histone deacetylase related to the yeast transcriptional regulator Rpd3p. Science, 272, 408–411.

Thul PJ, Åkesson L, Wiking M, Mahdessian D, Geladaki A, Ait Blal H, Alm T, Asplund A, Björk L, Breckels LM, Bäckström A, Danielsson F, Fagerberg L, Fall J, Gatto L, Gnann C, Hober S, Hjelmare M, Johansson F, Lee S, Lindskog C, Mulder J, Mulvey CM, Nilsson P, Oksvold P, Rockberg J, Schutten R, Schwenk JM, Sivertsson Å, Sjöstedt E, Skogs M, Stadler C, Sullivan DP, Tegel H, Winsnes C, Zhang C, Zwahlen M, Mardinoglu A, Pontén F, von Feilitzen K, Lilley KS, Uhlén M, Lundberg E (2017) A subcellular map of the human proteome. Science, 356, eaal3321.

Verhey KJ, Gaertig J (2007) The tubulin code. Cell Cycle, 6, 2152–2160.

Wu S, Rice JC (2011) A new regulator of the cell cycle: the PR-Set7 histone methyltransferase. Cell Cycle, 10, 68–72.

Yoo BK, Emdad L, Gredler R, Fuller C, Dumur CI, Jones KH, Cook-Jackson C, Su Z, Chen D, Saxena UH, Hansen U, Fisher PB, Sarkar D (2010) Transcription factor Late SV40 Factor (LSF) functions as an oncogene in hepatocellular carcinoma. Proc Natl Acad Sci U S A, 107, 8357–8362.

Zhang X, Sun F, Qiao Y, Zheng W, Liu Y, Chen Y, Wu Q, Liu X, Zhu G, Chen Y, Yu Y, Pan Q, Wang J (2017) TFCP2 Is Required for YAP-Dependent Transcription to Stimulate Liver Malignancy. Cell Rep, 21, 1227–1239.

Ziegelbauer J, Shan B, Yager D, Larabell C, Hoffmann B, Tjian R (2001) Transcription factor MIZ-1 is regulated via microtubule association. Mol Cell, 8, 339–349.

