## Supplemental Figures for "Transcription factor LSF facilitiates lysine methylation of α-tubulin by microtubule-associated SET8"

**Table S2. List of $\alpha$-tubulin peptides.**

| Peptide ID | Sequence |
|---|---|
| K40 | H-DGQMPSD**K**TIGGGDD-NH2 |
| K40-Ac | H-DGQMPSD**K-Ac**TIGGGDD-NH2 |
| K304 | H-PANQMV**K**CDPRHG-NH2 |
| K311 | H-CDPRHG**K**YMACCL-NH2 |
| K311A | H-CDPRHG**A**YMACCL-NH2 |
| K311S | H-CDPRHG**S**YMACCL-NH2 |
| K311-Me | H-CDPRHG**K-Me**YMACCL-NH2 |
| K311-Ac | H-CDPRHG**K-Ac**YMACCL-NH2 |

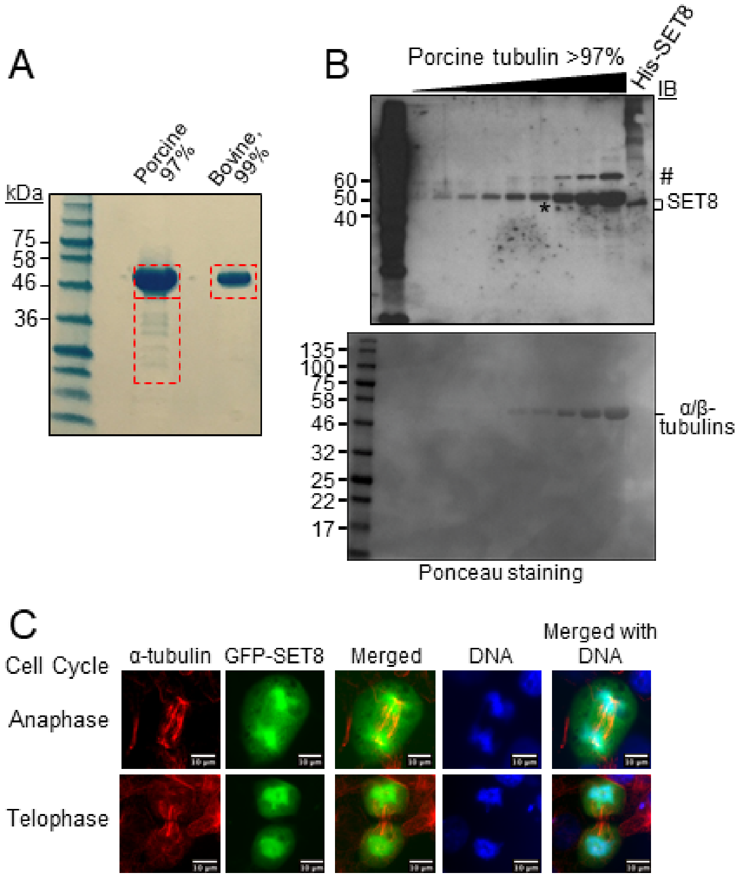

**Figure S1. Interactions between SET8 and α-tubulin.**

**A** Commercial, purified tubulin preparations from the indicated two sources were electrophoresed through a Tris-glycine gel, and the regions of the gels indicated by the dotted red boxes were excised for independent mass spectroscopy analysis. The results from the indicated gel slices of both tubulin preparations are presented in Appendix Table S1.

**B** Immunoblotting (IB) of purified porcine brain tubulin (rPeptide, >97%) using SET8 polyclonal antibody (Active Motif) shows a weak band (*) co-migrating with purified His-SET8. Top: Immunoblot, using biotinylated protein molecular weight markers. The most prominent band in the immunoblot, immediately above SET8, reflects nonspecific interaction of the antibody with the vast amount of α/β-tubulins; # indicates another nonspecific interacting protein. Bottom: Ponceau staining of same gel (shown in grayscale), using standard molecular weight markers. Due to the low levels of SET8 in the tubulin preparation, accurate quantitation of the relative levels of SET8 *versus* tubulins was not possible.

**C** Colocalization of SET8 and α-tubulin in COS7 cells at late mitotic phases. GFP-SET8 (green) was expressed in asynchronous cells, tubulin was detected with anti-α-tubulin antibody (red), and DNA with DAPI (blue). Yellow in the merged image indicates colocalization of SET8 and α-tubulin. Images are from cells identified as being in the indicated stages of cell cycle progression. These results were obtained as part of the same experiment shown in Fig 1A.

# A

Tubulin alpha chain (P02550)

**CONTROL:** 93% sequence coverage

| Confidence | Sequence | Modifications | # PSMs | Theoretical MH+ [Da] | Cross correlation (Xcorr) | Percolator q-value | Amino Acid Residues |
|---|---|---|---|---|---|---|---|
| High | AYHEQLSVAEITNACFEPANQMVK | 1×CAM [C15]; 1×Methyl [K24] | 1 | 2764.30691 | 5.18 | 0 | 281-308 |
| High | AYHEQLSVAEITNACFEPANQMVK | 1×OX [M22]; 1×CAM [C15]; 1×Methyl [K24] | 3 | 2780.30183 | 4.49 | 0.000487 | 281-308 |

**PLUS SET8:** 80% sequence coverage

| Confidence | Sequence | Modifications | # PSMs | Theoretical MH+ [Da] | Cross correlation (Xcorr) | Percolator q-value | Amino Acid Residues |
|---|---|---|---|---|---|---|---|
| High | AYHEQLSVAEITNACFEPANQMVKCDPR | 2×CAM [C15; C25]; 1×Methyl [K24] | 1 | 3292.51838 | 4.36 | 0 | 281-308 |
| High | HGKYMACCLLYR | 1×OX [M5]; 2×CAM [C7; C8]; 1×Methyl [K3] | 2 | 1601.73854 | 4.46 | 0 | 309-320 |
| High | YPRAHFPLATYAPVISAEKAYHEQLSVAEITNACFEPANQMVK | 1×OX [M41]; 1×CAM [C34]; 1×Methyl [K19] | 1 | 4892.41747 | 1.83 | 0.00136 | 262-304 |
| High | AYHEQLSVAEITNACFEPANQMVK | 1×OX [M22]; 1×CAM [C15]; 1×Methyl [K24] | 1 | 2780.30183 | 3.98 | 0 | 281-304 |
| High | TIQFVDWCPTGFK | 1×CAM [C8]; 1×Methyl [K13] | 1 | 1612.78283 | 2.08 | 0.00287 | 340-352 |
| High | AYHEQLSVAEITNACFEPANQMVK | 1×CAM [C15]; 1×Methyl [K24] | 1 | 2764.30691 | 5.05 | 0.000473 | 281-304 |

K304 in α-tubulin Endogenous α/β-tubulin

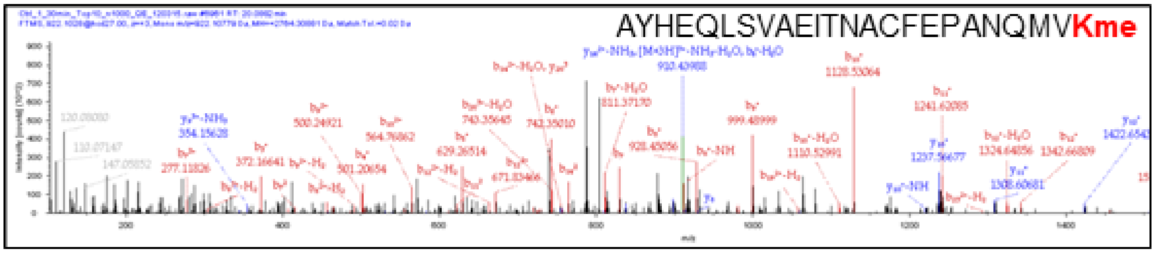

K280 in α-tubulin Endogenous α/β-tubulin plus SET8

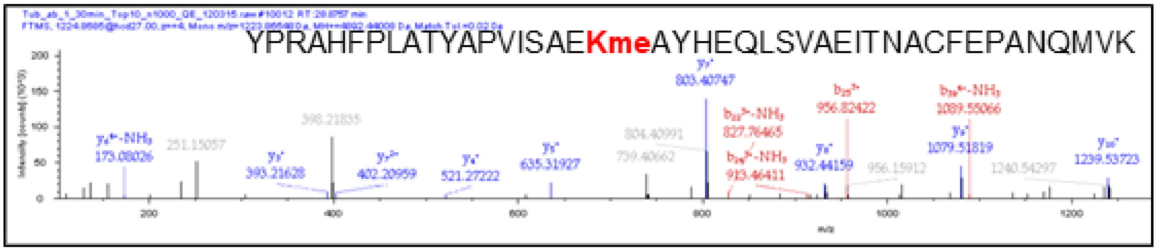

K352me in α-tubulin Endogenous α/β-tubulin plus SET8

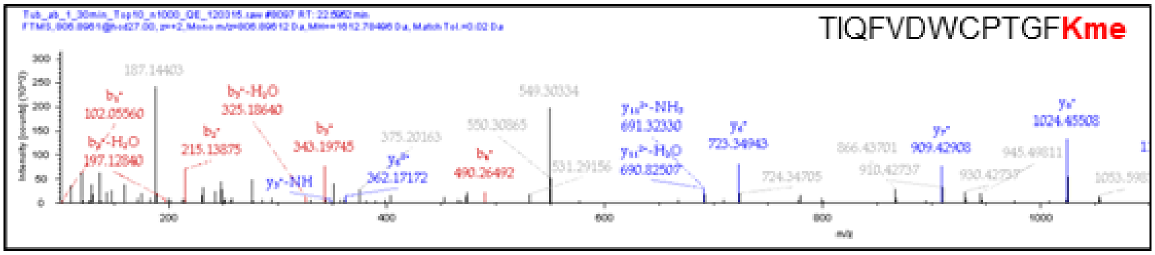

**B**

Tubulin beta chain (P02554)

**CONTROL**: 91% sequence coverage

| Confidence | Sequence | Modifications | # PSMs | Theoretical MH+ [Da] | Cross corre-lation (Xcorr) | Percolator q-value | Amino Acid Residues |
|---|---|---|---|---|---|---|---|
| High | MREIVHLQAGQCGNQIGAK | 1×CAM [C12]; 1×Methyl [K19] | 4 | 2124.0801 | 4.96 | 0 | 1-19 |
| High | ALTVPELTQQMFDAKNMMAACDPR | 1×CAM [C21]; 1×Methyl [K15] | 2 | 2752.2925 | 2.98 | 0 | 283-306 |

K19 in β-tubulin

Endogenoous α/β-tubulin

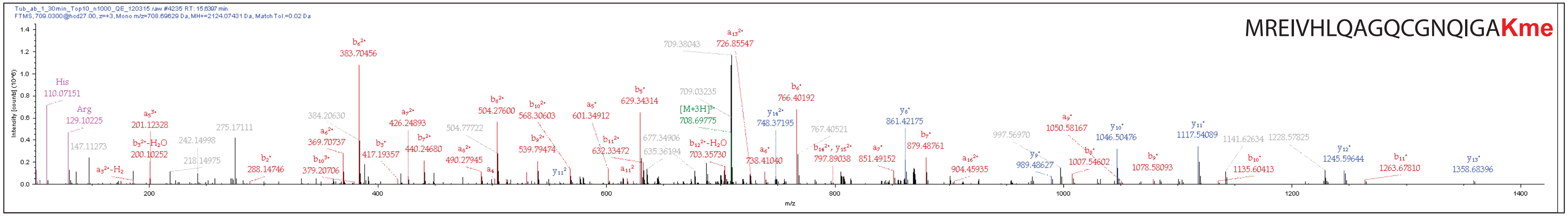

K297 in β-tubulin

Endogenoous α/β-tubulin

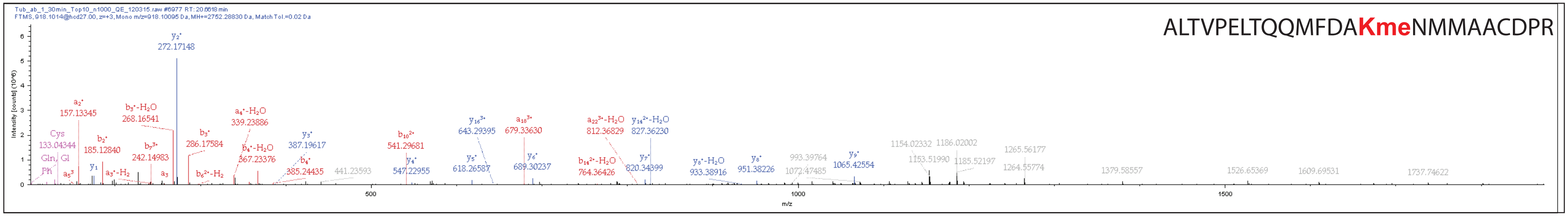

C

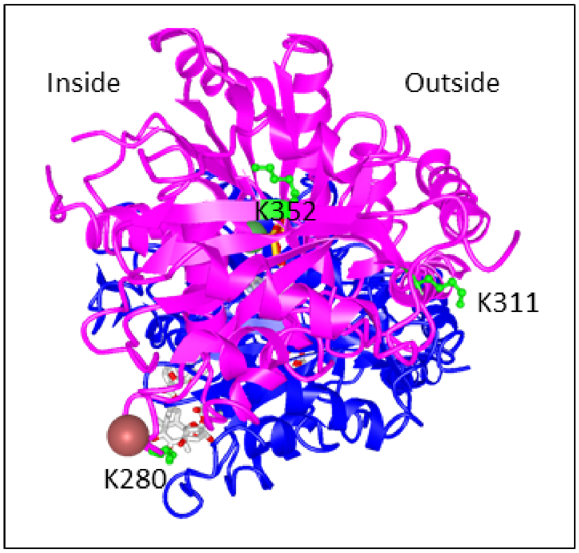

D

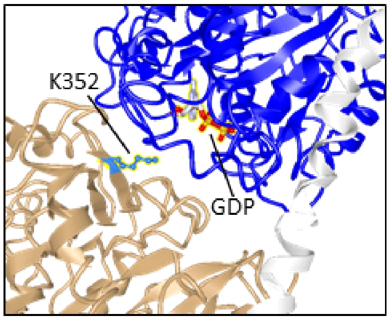

**Figure S2. Spectra and structural locations of specific methylated peptides in α-tubulin.**

**A** Top: statistics indicating identification of methylated lysine residues in α-tubulin by mass spectroscopy, observed both before and after incubation with SET8. Bottom: specific spectra demonstrating methylation of α-tubulin on K304 in tubulin purified from post-mitotic cells, plus methylation at K280 and K352 mediated by SET8 *in vitro*.

**B** Top: statistics indicating identification of methylated lysine residues in β-tubulin by mass spectroscopy. Bottom: specific spectra demonstrating methylation of β-tubulin on K19 and K297 in tubulin purified from post-mitotic cells.

**C** The 3-dimensional structure of the α/β-tubulin heterodimer (PDB ID 1JFF; purple: α-tubulin; blue: β-tubulin) from a top view, indicating positions of lysines targeted by SET8 (green). Notably, K352 is located on the upper surface of α-tubulin, K311 is located on the outside surface of the MT, and K280 is located toward the lumen (inside).

**D** In a matrix structure of multiple α/β-tubulin heterodimers (stabilized by stathmin; PDB ID 1FFX; brown: α-tubulin; blue: β-tubulin; light grey: stathmin), K352 (green) is closely located near the GDP binding site in β-tubulin. Based on the position of K352 at the α/β-tubulin interface, K352 methylation would be likely to impact tubulin polymerization. In contrast, modification of K311 does not suggest direct consequences to tubulin polymerization, although by altering which proteins bind the MTs it could modify polymerization dynamics.

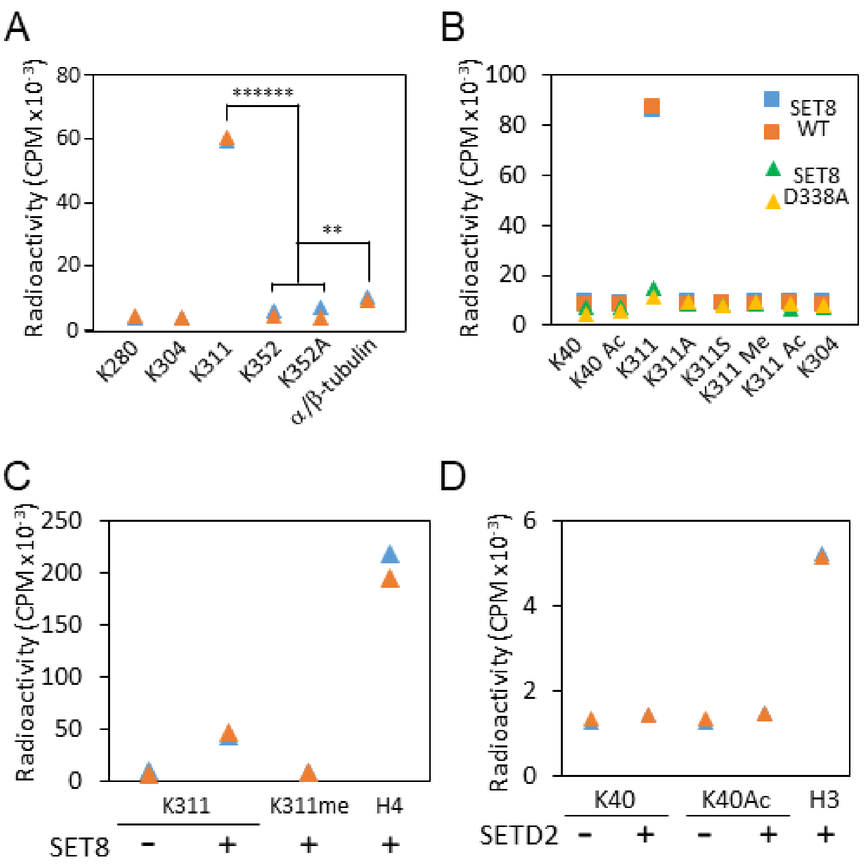

**Figure S3. K311 of α-tubulin is methylated by SET8 *in vitro*.**

**A** *In vitro* peptide methylation assay of candidate α-tubulin methylation sites (see Appendix Table S2), mediated by purified GST-SET8. The K311-containing peptide, but not the other peptides, is methylated to a statistically significant extent (****** $p = 6x10^{-7;}$ standard one-tailed, equal variance t test). This peptide demonstrates much higher radioactive incorporation compared to α/β-tubulin, due to addition of equal µg of the substrates, and therefore a 69-fold higher molar concentration of the target lysine in the peptide reaction, compared to the total protein reaction. Nonetheless, α/β-tubulin was also methylated to a statistically significant extent (** p = 0.009, standard one-tailed, equal variance t test), compared to level of radioactivity detected in the peptide containing K352 and its negative control K352A.

**B** *In vitro* peptide methylation assay of α-tubulin K40-, K311-, and K304-containing peptides of wild type or mutant sequences (see Appendix Table S2 for peptide sequences). Peptides were incubated with either SET8 WT (active enzyme) or SET8 D338A (inactive enzyme). Only the unmodified K311-containing α-tubulin peptide was specifically methylated.

**C** Comparision of the methylation *in vitro* of the α-tubulin K311-containing peptide and histone H4 by GST-SET8. Note that scale is different from those in panels **A** and **B**.

**D** A parallel *in vitro* methylation reaction by SETD2 of its known target, histone H3 (note the 40-fold more sensitive scale), as well as a α-tubulin K40-containing peptide.

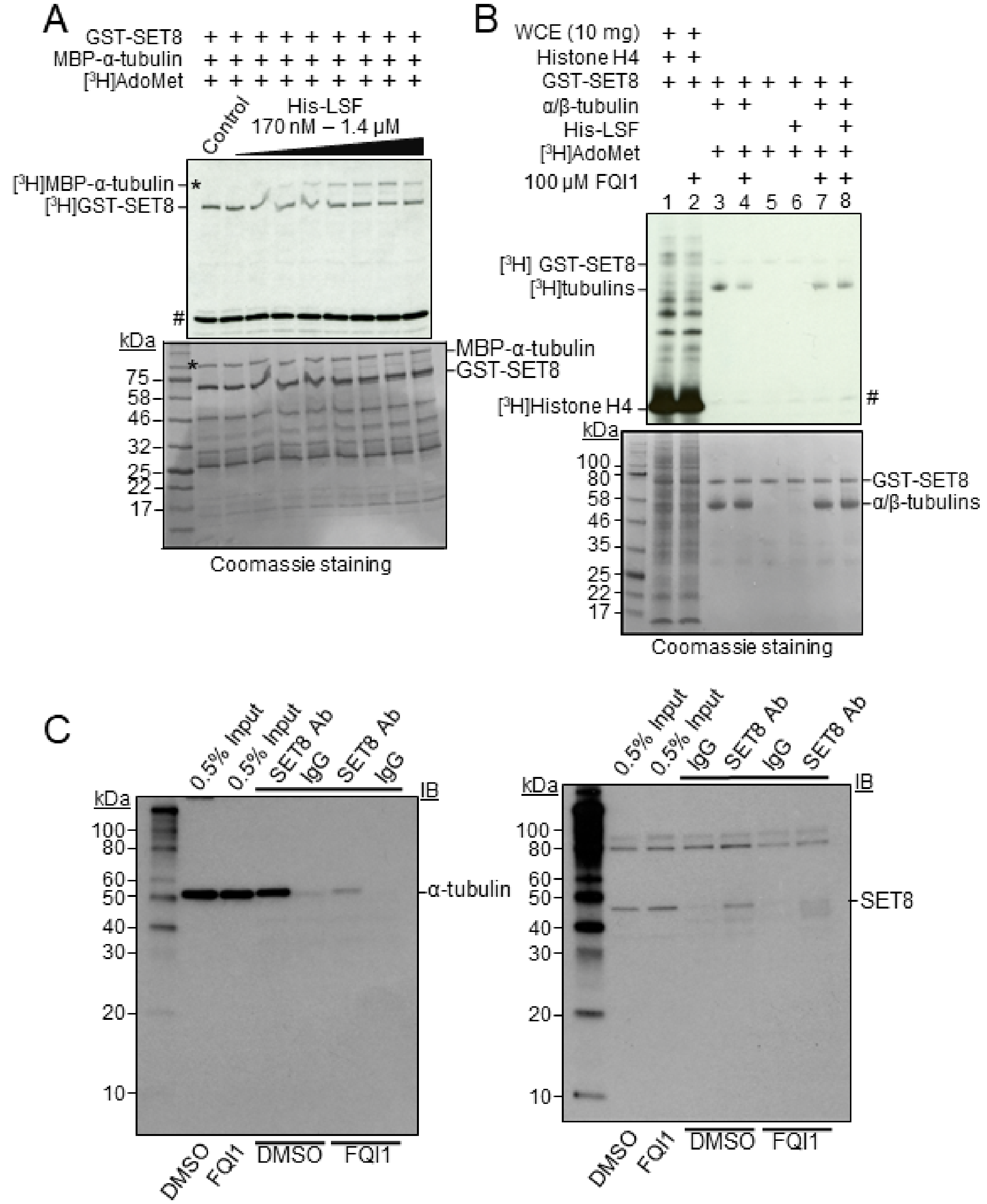

**Figure S4. LSF and FQI1 modulate α-tubulin methylation by SET8.**

**A** Recombinant MBP-α-tubulin methlyation reactions were performed with addition of the indicated, increasing range of concentrations of His-LSF. Top: autoradiogram of methyltransferase assays showing the methylated MBP-α-tubulin (*), plus automethylation of GST-SET8. The higher relative levels of GST-SET8 to MBP-α-tubulin in this experiment led to greater initial automethylation of SET8 relative to tubulin methylation. # indicates the migration of [$^3$H]-labeled impurities. Bottom: Coomassie-stained gel (shown in grayscale) showing relative levels of protein components. Protein bands <50 kDa are protein degradation products in the GST-SET8 preparation, which are particularly evident in this experiment.

**B** FQI1 prevents LSF-mediated increase in methylation of tubulin by SET8. Top: autoradiogram of methyltransferase assays with purified tubulin (>99%, MP-Biomedical) in the presence of FQI1 or vehicle (DMSO), and with or without 862 nM His-LSF.
WCE: whole cell extract, as a control for methylation of histone H4 and other proteins.
### indicates the migration of [$^3$H]-labeled impurities. Bottom: Coomassie-stained gel (shown in grayscale) showing relative levels of protein components.

**C** Co-immunoprecipitation of endogenous α-tubulin (left: immunoblotted with anti-α-tubulin from Sigma T9026) with endogenous SET8 (using anti-SET8 from Active Motif for the immunoprecipitation) was analyzed in HEK293 cell lysates following treatment of the cells with 2.5 μM FQI1 or vehicle for 24 hr. The level of immunoprecipitated endogenous SET8 (right: immunoblotted with anti-SET8 from Santa Cruz), although also reduced in this experiment upon treatment with FQI1, was diminished to a lesser degree than that of α-tubulin.
